# Head before heart? Cognitive empathy may emerge before affective empathy in the developing brain

**DOI:** 10.1101/2025.03.24.645025

**Authors:** C. Bulgarelli, P. Pinti, T. Bazelmans, A. F. de C. Hamilton, E.J.H. Jones

## Abstract

Empathy is crucial for social interactions across all cultures, and is foundational to establishing social cooperation and group ties in human societies. Challenging the current predominant view, we recently proposed that understanding others’ emotions (cognitive empathy) might emerge earlier than actually sharing those emotions (affective empathy) (Bulgarelli & Jones, 2023). Here we test this hypothesis by measuring which empathic component matures first during toddlerhood, a critical period for the development of broader social networks. Addressing this question is critical to understand the mechanisms through which caregivers scaffold empathy development. Traditional approaches are inadequate, as they rely on children’s verbal skills or unfamiliar scenarios that lack ecological validity.

In this preregistered study, we employed a novel toddler-appropriate task to dissociate neural and physiological correlates of cognitive and affective empathy in N=90 3-to-6-year-olds using functional near-infrared spectroscopy (fNIRS) and simultaneous heart rate monitoring to identify internal markers of empathy.

We found that brain regions supporting affective and cognitive empathy in young children resemble those observed in adults. Importantly, we showed an effect of age on network specialisation with brain activations of cognitive empathy stronger in younger compared to older preschoolers, and brain activations of affective empathy stronger in older compared to younger preschoolers. These results may provide the first evidence that cognitive empathy is engaged early and develops alongside or ahead of affective empathy in preschoolers, challenging existing models and suggesting a new framework for understanding the development of empathy.

**Significance statement:** Empathy is a crucial social skill, consisting of an affective component—sharing emotion—and a cognitive component—conceptually understanding emotions. While to date it has been predominately accepted that the affective component develops earlier than the cognitive one, we recently proposed the opposite, which could have important implications for understanding the mechanisms that underpin social skills. Here we successfully developed a new task to dissociate physiological and neural markers of affective and cognitive empathy in N=90 preschoolers. We found that brain activations for cognitive empathy are stronger than for affective empathy in young children. This may provide the first direct evidence for the earlier emergence of cognitive empathy, suggesting that scaffolding the understanding of others’ emotions may be crucial for empathy development.

## Introduction

Sharing emotion is universal: a good friend is overjoyed when she gets promoted at work, we recognise why the news matters to her, and we genuinely feel happy for her. Or our sister loses her beloved pet, we understand what the loss means to her, and we feel sad for her loss. This ability to share someone else’s happiness or sadness and understand the reason for their emotion is called empathy, and it pervades and scaffolds our everyday social interactions. Empathy is a social skill crucial for successful social interactions (1). This ability is present across cultures (2) and is foundational to establishing social cooperation and group ties in human societies (3, 4). Importantly, empathy is believed to play a crucial role in the evolutionary development of humans as a social species (5), with evidence of this skills also observed in primates (6), suggesting its pivotal role in our society from both ontogenetic and phylogenetic perspectives (7).

Empathy consists of two components: an affective dimension, which involves sharing others’ emotions, and a cognitive dimension, which involves conceptually understanding others’ emotions. Interestingly, several neuroimaging studies on adults documented that these two components are supported by different brain networks (8–10), whose functional roles are partly overlapping (11). Cognitive empathy has been consistently linked to the dorsolateral prefrontal cortex (DLPFC) and the temporoparietal junction (TPJ) (8, 12–14). Affective empathy has been associated with activity in medial prefrontal cortex (MPFC) and superior temporal gyrus (STG). Notably, the MPFC has also been associated with cognitive aspects of empathy (15, 16). Cognitive empathy can involve elements of theory of mind or mentalising (5) and activates similar regions in adults (14, 17), but specifically refers to understanding others’ emotional states rather than the broader ability to represent another person’s thoughts (4, 18). Thus, cognitive empathy may not necessarily *require* the broader belief-representing capacities of theory of mind, as emotional states can be read from external cues.

While empirical studies on empathy are abundant in adults, we know little about the development of this fundamental social skill during the first years of life. In particular, the developmental ordering of its affective and cognitive components, and the neural changes underlying it, remain debated. However, understanding the mechanisms behind the emergence of the two empathic components is crucial for advancing our understanding of social development and the factors that can support emerging bonds for children in their natural environments. So far, it has been widely accepted that the affective component of empathy emerges earlier than the cognitive one. In a prior theoretical paper (19), we termed this *foundational affective empathy model*, in which emotional contagion in early infancy - an automatic sharing of emotions and behaviours without a conscious understanding of another’s emotional state - develops into affective empathy (7). This is thought to then provide the foundation for children learning to label or conceptually identify other people’s emotions based on their own sharing of these emotions - cognitive empathy. Recently, this linear, stage-based account has been challenged by evidence that affective and cognitive dimensions of empathy appear to be present together as early as the first year of life (19–23), which is difficult to reconcile with a sequential stage account in which one component gives rise to the other. At the same time, much of the earlier evidence taken to support an affective-first developmental sequence relied on a small corpus of studies, mostly exploring children’s behavioural or physiological reaction to the experimenter in pain (24) or a doll crying (25). However, these tasks do not reflect typical daily life situations, and do not consider the extensive body of literature suggesting that we tend to experience greater empathy for individuals we perceive as more similar to ourselves (26), which would be another child when assessing empathy in children. Additionally, these tasks primarily investigated empathy in response to negatively connoted emotional events, such as exposing children to someone in pain or distress. These stimuli may directly cause distress in the child watching (just like any other loud environmental sound), rather than the child necessarily feeling a shared emotion (being sad *for* someone else, not *because* of someone else or for yourself). “Negativity bias,” refers to the tendency to exhibit stronger reactions to negative emotions and events than to positive ones (27). While it is plausible to think that initial empathic responses first emerge for others’ negative emotions (e.g., sadness, fear, anger) and later extend to a broader range of emotions, to substantiate this claim, we need tasks that elicit empathy for both positively and negatively connoted emotional events. Within the *foundational affective empathy model*(*19*), cognitive empathy does not emerge before the age of 4. This has often been based on tasks that rely on children’s verbal abilities (28, 29), but it confounds the concept of cognitive empathy with a child’s ability to describe their inner experiences. Lastly, most studies have explored affective and cognitive empathy using different sets of stimuli for the two components (for example see 12). However, since these are two aspects of the same multifaceted skill, the same stimuli must be used in empirical tasks to elicit empathy and measure both its affective and cognitive components.

In contrast to the prevailing theoretical model, lived experience indicates that parents and caregivers spend a considerable amount of time labelling and describing other people’s emotions to young children. Observations in nurseries indicate that the predominant response of young children to another child crying is either ignoring them or watchful interest, rather than any display of a matched emotion (19). Nursery staff will commonly follow by labelling and explaining the child’s emotional state (“Look, Bobby is sad because he dropped his toy!”). This indicates a rich context for children to develop cognitive understanding, but also suggests that by feeling the need to explain, caregivers do not operate on the belief that toddlers intuitively respond to another toddler’s emotions by feeling the same emotion themselves (which may be more likely to elicit comments like “Ahh, you are feeling sad for Bobby”). Coupled with the lack of convincing empirical evidence for foundational affective empathy, we proposed a *foundational cognitive empathy model*, in which a conceptual understanding others’ emotions is at the basis of the experience of empathy (19). Consistent with this, it has been shown that children younger than 4 years of age have some understanding of others’ emotions (though still assessed with a verbal emotion labelling task) (30) and that they actively try to understand others’ condition when they see them in distress (31). The pivotal scaffolding role of significant adults in childcare environments might be crucial for empathy development, as it is for other social skills. It has been shown for example that parents’ mind-mindness and mental state language predicted their children’s theory of mind and emotion understanding (32–34). It is therefore plausible to think that by observing how teachers or parents react to another child in distress, children learn what it means for their classmate to feel sad, even if the visceral experience of shared sadness has not yet developed. This framework is consistent with what we observed during 21 hours of naturalistic observations in a nursery class, where several episodes of preschoolers approaching and paying attention to a classmate crying occurred after the teacher approached the child in distress (19).

To directly evaluate the explanatory value of the two frameworks we proposed in prior theoretical work (19) - the *foundational cognitive empathy model* against the *foundational affective empathy model* - we designed a novel task to comparably assess both affective and cognitive empathy in preschoolers. The preschool period is a critical window for assessing empathy development. After the second year of life, children are capable of separating emotions in others from emotions in themselves, a critical prerequisite for empathy (35, 36). Additionally, the vast majority of children older than 3 attend some form of daycare (at least in the UK where this is partially government-funded), where they begin to engage more intensively in social interactions with their peers and thus where empathy becomes increasingly relevant to everyday interactions. Addressing the limitations of tasks used in previous research, the key characteristics of our task included: i) the presentation of scenarios featuring preschoolers as characters, and both positively and negatively connoted emotional events; ii) the simultaneous assessment of affective and cognitive empathy using the same stimuli; iii) the decision to avoid exclusively relying on verbal skills to evaluate cognitive empathy. To go beyond the preschoolers’ behavioural and verbal manifestations of empathy, during the task we recorded neural activations supporting affective and cognitive empathy using functional near-infrared spectroscopy (fNIRS), a child-friendly neuroimaging method that uses near-infrared light to measure changes in oxygenated (HbO_2_) and deoxygenated blood (HHb) in the brain, as a proxy for brain activation (37). fNIRS has proven to be an excellent tool for neuroimaging studies with developmental populations; it is highly tolerant to motion, allows for the assessment of awake participants, enabling the investigation of social skills for example, and is well accepted by young participants (38).

Building on our theoretical model (19) , we preregistered our hypotheses regarding brain activations for this novel task (https://osf.io/crvz2). We predicted that preschoolers would show:

i) greater neural activations of the frontal cortex and the temporal lobe for the emotional rather than neutral scenarios, as a neural marker of empathy (H1). While previous studies have investigated neural responses to others’ pain or distress as the empathy-eliciting stimulus, which may evoke personal distress in the observing child rather than a shared emotional response, and have largely been restricted to negatively valenced events (39–41), this is the first study to test whether the developing preschool brain responds to emotional scenarios in regions comparable to those documented in adults, using a task that assesses both empathy components with the same stimuli.
ii) greater neural activations of the frontal cortex and the temporal lobe for the negative rather than positive emotional scenarios, especially in younger toddlers (H2), which would be consistent with the “negativity bias” hypothesis (27). This is the first study to investigate brain activations in response to both positive and negative emotional events in young children.
iii) developmental changes, and in particular stronger neural markers of cognitive empathy in younger compared to older preschoolers and stronger neural markers of affective empathy in older compared to younger preschoolers (H3). This would provide the first empirical evidence supporting our proposed *foundational cognitive empathy model*, challenging existing frameworks and revolutionising our understanding of empathy development.

As adult studies have shown that arousal changes during empathic reactions, especially in response to negative emotions (42), during the task simultaneously to fNIRS we also recorded preschoolers’ heart rate through electrocardiogram (ECG) recordings as a physiological indicator of arousal. Our preregistered hypothesis was that preschoolers would show larger change in heart rate between the set-up of the scene and the event during emotional scenarios, particularly those involving negative emotions, compared to neutral ones. Figure 1 shows a graphical representation of the hypotheses of this work based on our task design.

**Figure 1.**
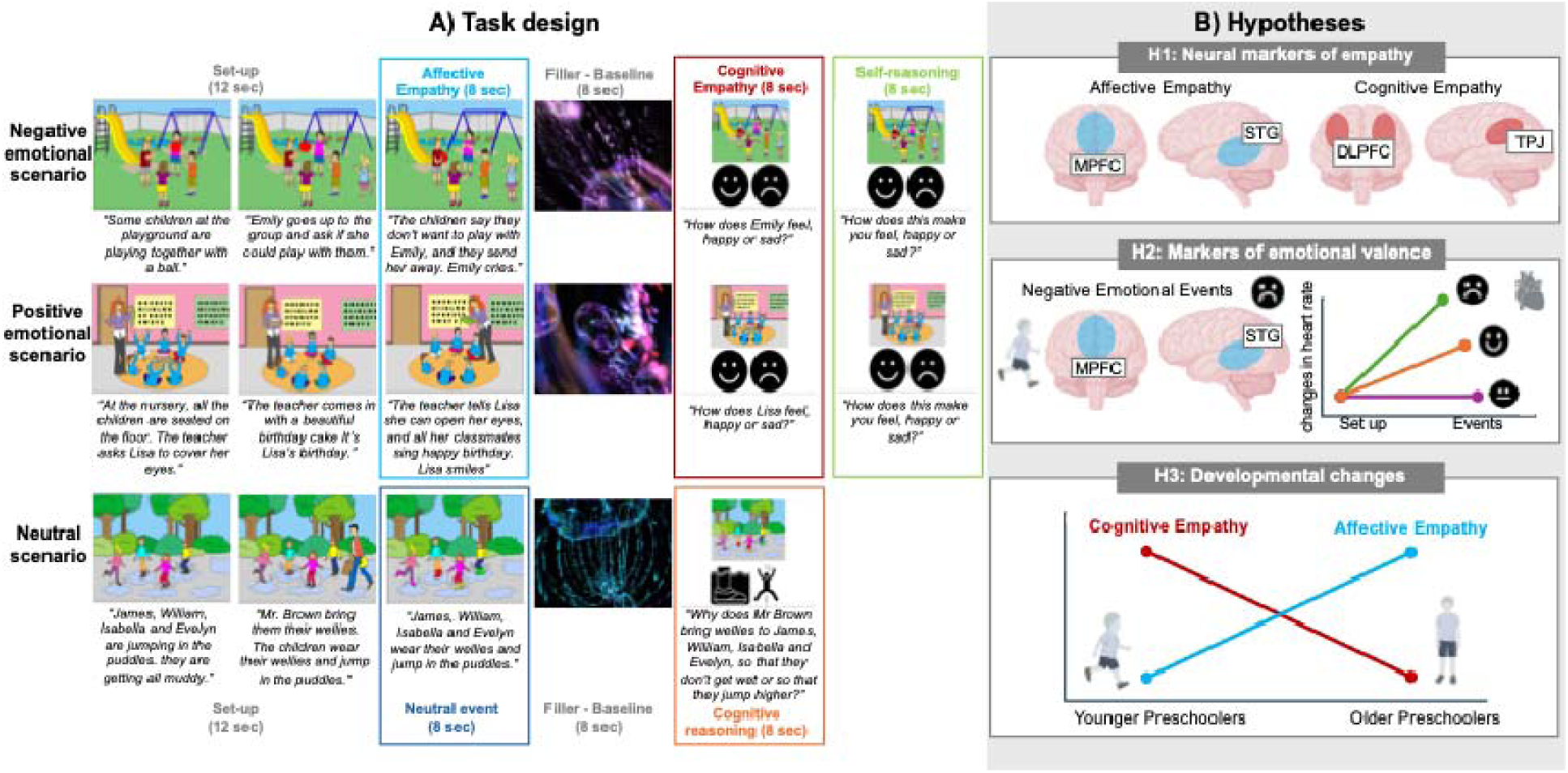
A) Task design. Participants saw 8 emotional scenarios (4 negative and 4 positive) where each trial has an affective empathy phase, followed by a short filler and then a cognitive empathy question and a self-reasoning question. Neutral trials followed the same structure with neutral scenarios. B) Hypotheses (all pre-registered). H1: Affective empathy would engage MPFC and STG, while cognitive empathy would engage DLPFC and TPJ. H2: Negative scenarios should lead to greater engagement of affective empathy brain regions (especially in younger preschoolers) and greater changes in heart rate between set-up and events of the scenarios compared to positive or neutral. H3: Developmental changes, with neural markers of cognitive empathy preceding affective empathy, in line with the *foundational cognitive empathy model*. Icons of this figures were made using Biorender.com (https://www.biorender.com/).

## Results

N = 95 preschoolers watched 16 scenarios (8 emotional, with 4 positive and 4 negative, and 8 neutral) featuring child characters experiencing emotional or neutral events, while neural activations and heart rate were recorded. In emotional scenarios, the emotional event served to elicit affective empathy, and was followed by a question about the character’s emotional state (’how does s/he feel?’, cognitive empathy), a question about the child’s own feelings (’how does this make you feel?’). Neutral scenarios followed the same structure with a non-emotional event and a factual question. Full details are provided in the Methods (see also Figure 1A).

### Verbal responses to the task questions

First, we assessed compliance with our task by evaluating how many of the 16 proposed trials (8 emotional and 8 neutral) participants attended (out of the total N=95 of participants who observed the stimuli). 88.42% (N= 84) of participants completed all 16 trials (mean number of emotional scenarios attended = 7.83/8, mean number of neutral scenarios attended = 7.79/8). Among those who did not complete all trials, all attended more than 75% of the emotional scenarios, while 63.64% (N=7) attended more than 75% of the neutral scenarios. There was no significant effect of age on the number of trials included (all p > 0.1). This suggests that our novel task is highly suitable for children aged 3 to 5 years.

We then explored verbal responses to the task questions, both in the emotional and neutral scenarios. 86.32% (N=82) of the participants answered congruently with the character’s emotions at the first emotional question (‘*how does s/he feel?*’), 72.63% (N=69) of the participants answered congruently (i.e. what we would expect the child to feel) with the character’s emotions at the second emotional question (‘*how does this make you feel?*’), and 55.79% (N=53) of the participants made no mistakes, and 26.32% (N=25) made one mistake on the question in the neutral scenarios. Older age was associated with a lower percentage of mistakes when asked how the other child felt in the emotional scenarios (*r*(93) = -0.34, *p* = 0.001) and about events within the neutral scenarios (*r*(93) = - 0.35, *p* = 0.001). Age did not associate with accuracy when asked ‘*how does this make you feel?*’ (*r*(93) = -0.15, *p* = 0.157).

### Neural markers of affective and cognitive empathy (H1)

To investigate which brain regions supported affective empathy in preschoolers, a one-sample t-test on the beta values comparing emotional negative and positive events with neutral events was conducted for each region of interest (ROI). Beta values index the strength of the haemodynamic response for each condition derived from a general linear model; a positive value reflects greater activation for the condition of interest relative to the comparison condition (see Methods). We found that preschoolers showed greater activations for the emotional compared to neutral scenarios in the left STG for both HbO_2_ (t(86) = 3.78, p < 0.001, q = 0.002, d = 0.41) and HHb (t(86) = 2.66, p = 0.005, q = 0.016, d = 0.29, both surviving FDR correction).

To investigate which brain regions supported cognitive empathy in preschoolers, a one-sample t-test on the contrast of beta values when participants were asked how the child felt in the emotional negative and positive scenarios versus the beta values for questions about facts in the neutral scenarios was conducted for each ROI. We found that preschoolers showed greater activations for the questions about the character’s emotional states compared to questions about neutral facts in the right MPFC (t(79) = 3.43, p < 0.001, q = 0.002, d = 0.38, surviving FDR correction), left DLPFC (t(79) = 3.41, p = 0.009, q = 0.020, d = 0.27, surviving FDR correction), right DLPFC (t(79) = 4.71, p < 0.001, q = 0.001 , d = 0.52, surviving FDR correction), right IPL (t(87) = 2.49, p = 0.007, q = 0.022, d = 0.26, surviving FDR correction) and right TPJ (t(86) = 1.95, p = 0.024, q = 0.051, d = 0.20) for HbO_2_ (no ROI showed significant different activation for HHb) (Fig.2).

**Figure 2.**
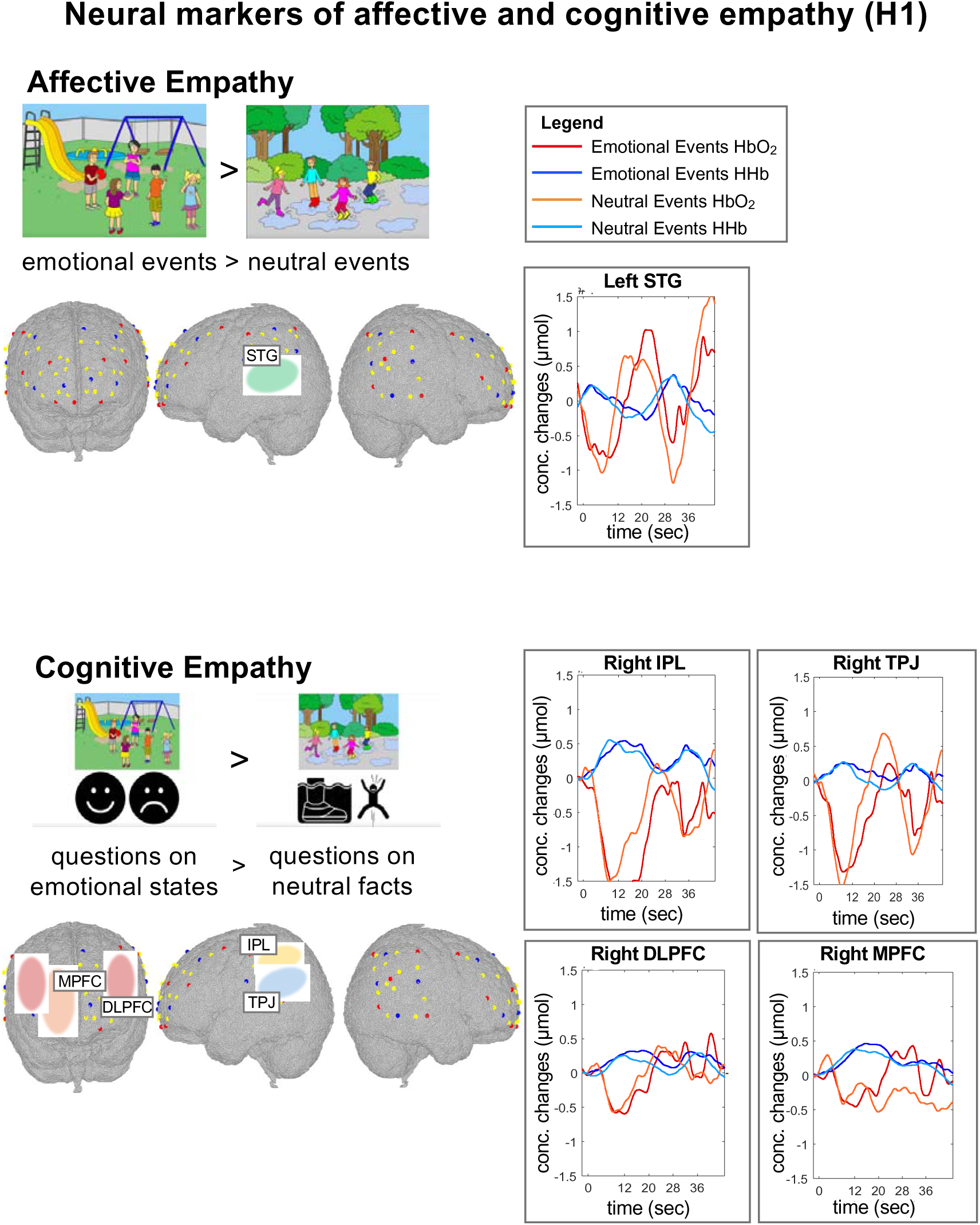
Brain regions supporting affective empathy (emotional > neutral events) and cognitive empathy (questions on emotional state > questions on neutral facts) and HRF plots. The images of the brains represent a schematic representation of the channels overlapped on a 5-year-old multilayer head model mesh. Regions are colour-coded as in Figure 4 in (31). The yellow band in the HRF plots represent the time window considered in the GLM analyses.

These results highlight that regions underpinning affective and cognitive empathy in preschoolers largely resemble those documented in adults.

### Neural markers of negative and positive emotional events (H2)

To investigate which brain regions showed greater activations for emotional negative rather than positive scenarios in preschoolers, a one-sample t-test on the contrast of beta values for emotional negative events versus the beta values for emotional positive events was conducted for each ROI. We found that preschoolers activated more left TPJ (t(87) = 2.46, p = 0.008, q = 0.020, d =0.26 , surviving FDR correction), right TPJ (t(86) = 2.92, p = 0.002, q = 0.010, d = 0.32, surviving FDR correction) and right STG (t(85) = 2.63, p = 0.005, q = 0.010, d = 0.28, surviving FDR correction) for the emotional negative compared to the emotional positive events, and left MPFC (t(79) = 3.64, p < 0.001, q = 0.002, d = 0.40 , surviving FDR correction) and right MPFC (t(79) = 2.09, p = 0.021, q = 0.042, d = 0.23, surviving FDR correction) for the emotional positive compared to the neutral events for HbO_2_ (no ROI showed significant different activation for HHb) (Fig.3).

**Figure 3.**
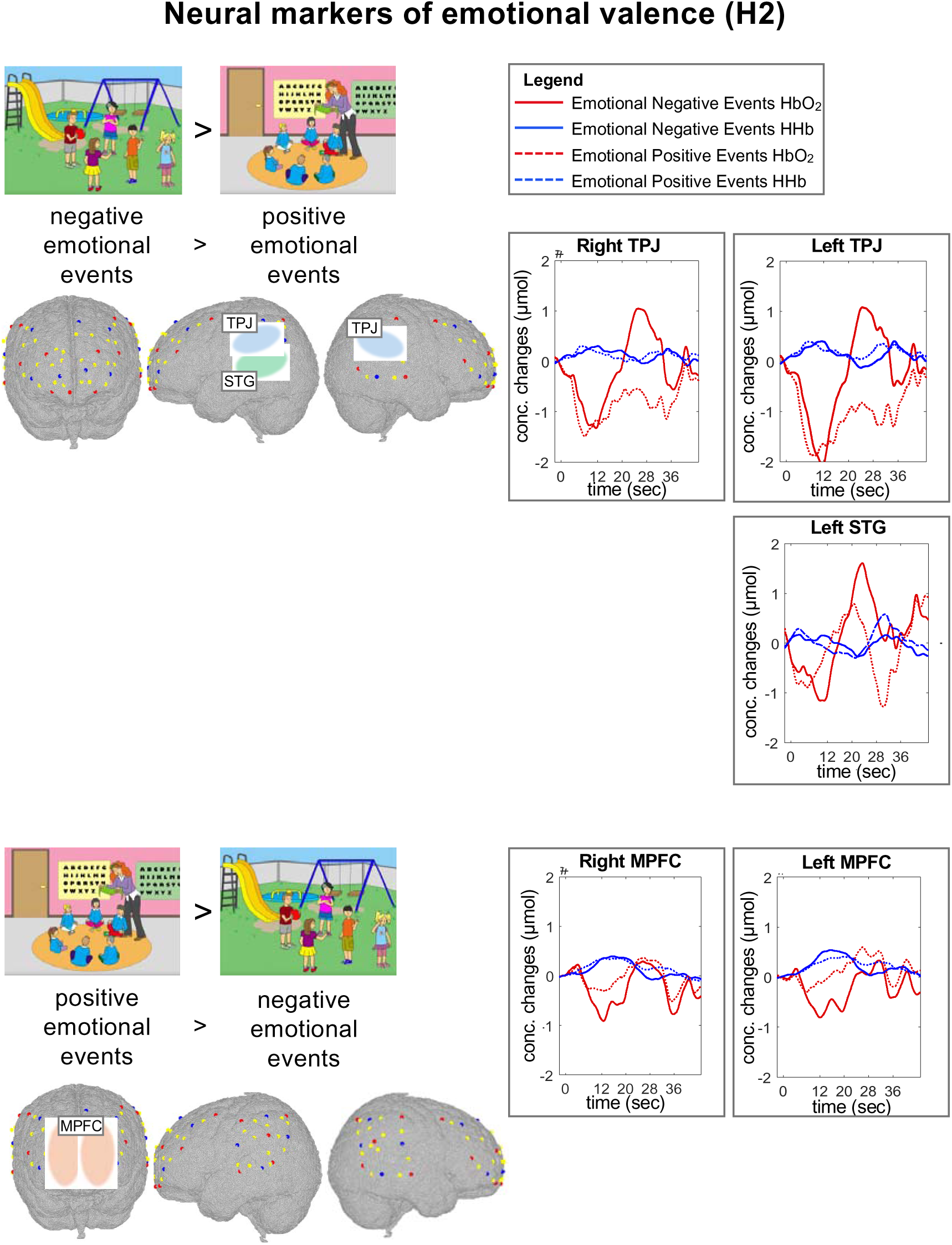
Brain regions for negatively and positively emotional scenarios and HRF plots. The images of the brains represent a schematic representation of the channels overlapped on a 5-year-old multilayer head model mesh. Regions are colour-coded as in Figure 4 in (31). The yellow band in the HRF plots represent the time window considered in the GLM analyses.

To investigate whether there was an association between age and brain activations for events with different emotional valences, we performed correlational analyses between age and brain regions that showed differences in activation between emotional negative and positive events. We found no significant associations (all p > 0.4). We additionally investigated the effect of age on brain activations for events with different emotional valences using age as a categorical variable (younger and older preschoolers, based on a median split at age 4.35 years, N=45 per age group). Independent t-tests between younger and older preschoolers on the regions that showed significant differences between the two emotional events at the group level showed no significant differences (all p > 0.2).

### Physiological markers of empathy and emotional valence: heart rate measures

To investigate whether there was a change in participants’ heart rate when attending emotional but not neutral events as hypothesised, we performed a repeated-measures ANOVA on the preschoolers’ average interbeat interval (IBI) with part of the scenario (set-up, event) and condition (emotional negative, emotional positive and neutral) as within-participant factors. We found a significant main effect of part of the scenario (*F*(1, 90) = 31.68, *p* < 0.001, η = 0.042), a significant main effect of condition (*F*(2, 90) = 27.22, *p* < 0.001, η = 0.078), and a significant interaction between part of the scenario and condition *(F*(2, 90) = 22.66, *p* < 0.001, ^2^ = 0.112) (Fig. 4A). Post-hoc paired t-η tests revealed an increase in the IBIs (i.e. greater IBI values, lower heart rate) between the event and set-up parts of the scenarios only in the emotional negative (*t*(90) = 7.52, *p* = 0.001) and emotional positive conditions (*t*(90) = 2.88, *p* = 0.005), but not in the neutral condition (*t*(90) = 0.65, *p* = 0.513). In addition, post-hoc paired t-tests revealed significantly longer IBIs (i.e. lower heart rate) in the emotional negative compared to emotional positive (*t*(90) = 6.90, *p* < 0.001) and neutral events (*t*(90) = 11.24, *p* < 0.001), and an almost significantly slower in the emotional positive compared to neutral events (*t*(90) = 1.93, *p* = 0.056).

**Figure 4.**
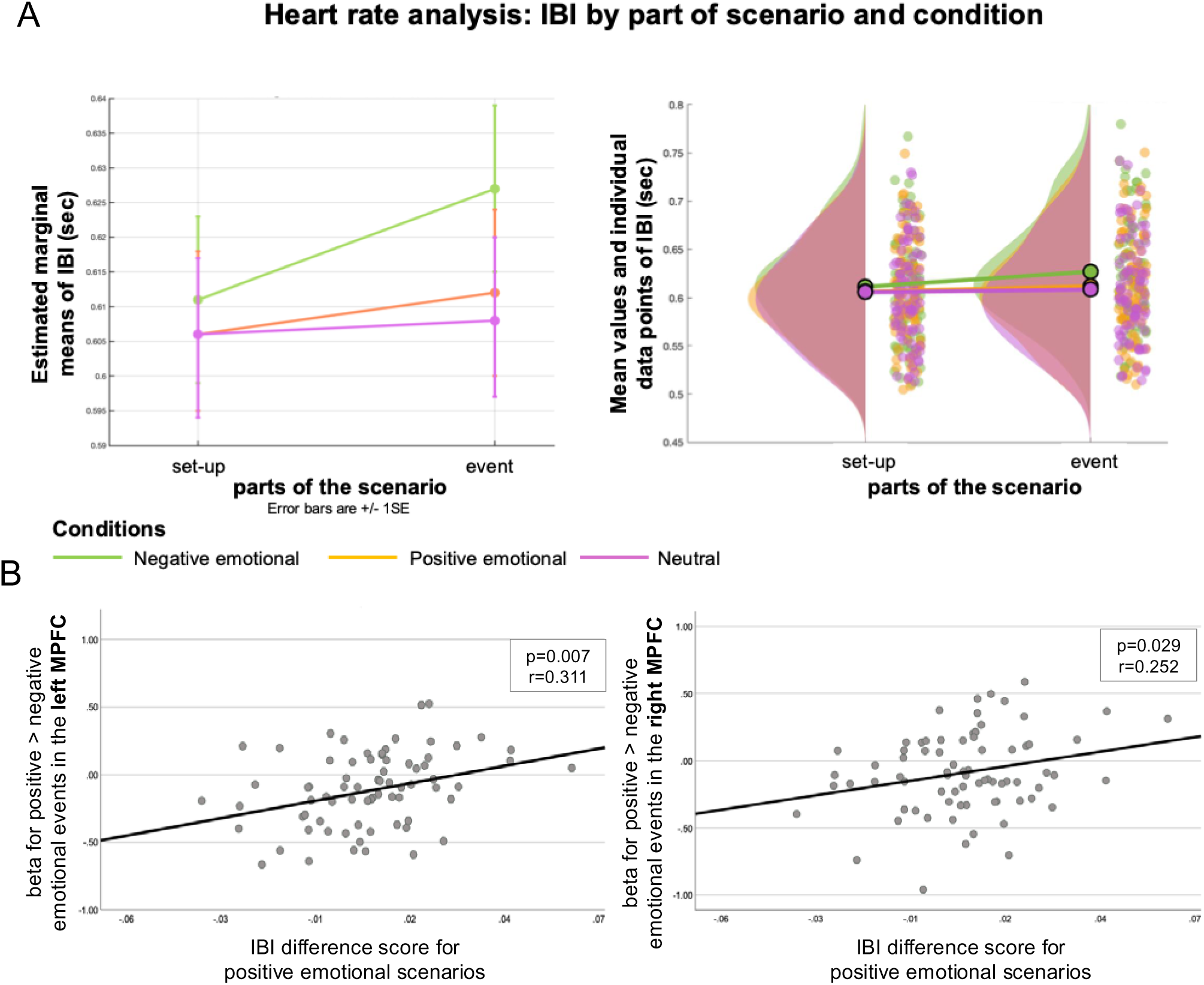
A) On the left, bar plots of the estimated marginal means of the IBI values per condition and part of the scenario. Error bars indicate +/- 1 SE. On the right, mean values and individual data points of the IBI values per condition and part of the scenario (created using the Raincloud Plots package (92)). B) Scatterplots representing the statistically significant relationships between the IBI difference score (event minus setup) for positive emotional scenarios and betas greater for positive compared to negative emotional relevant events in the left and right MPFC.

Although we did not hypothesise any specific association between physiological and neural underpinnings of affective empathy (emotional event vs neutral event), we exploratorily performed correlations between heart rate and brain activations underpinning affective empathy. We first calculated an IBI difference score by subtracting the IBI during the event phase of the scenarios from the IBI during the set-up phase of the scenarios for each condition. This IBI difference score was then used in correlational analyses with the beta values of the brain regions that showed significant activations in H2. There was a significant positive association between the IBI difference score for positive scenarios and greater activation of left and right MPFC for positive compared to negative events (left MPFC: *r*(75) = 0.311, *p* = 0.007, right MPFC: *r*(75) = 0.252, *p* = 0.029) (Fig. 4B).

### Networks specialisation increasing with age (H3)

To investigate whether age had any effect on the brain activations underlying affective and cognitive empathy, we performed ANCOVA analyses on the regions that showed significant differences in activation between the two empathy components (affective empathy, i.e., emotional events > neutral events, and cognitive empathy, i.e., questions about emotional states > questions about neutral facts), with age as a continuous covariate. We found that age had a significant effect on left STG for affective empathy (HbO_2_: *F*(1, 86) = 3.99, *p* < 0.049, η ^2^ = 0.045; HHb: *F*(1, 86) = 16.92, *p* <0.001, η*_p_*^2^ = 0.166) and an almost significant effect on right TPJ for cognitive empathy (HbO_2_: *F*(1, 86) = 3.60, *p* < 0.060, η*_p_*^2^ = 0.041). To explore the direction of the effect of age, we ran regression analyses. We found that age significantly predicted greater left STG activation supporting affective empathy, both for HbO_2_ and HHb (HbO_2_: B = 0.16, β = 0.21, t(86) = 2.00, p = 0.049; HHb: B = 0.09, β = 0.41, t(86) = 4.11, p < 0.001)). Age also negatively predicted greater right TPJ activation for the questions about emotional states compared to questions about neutral facts for HbO_2_, with younger age associated with greater right TPJ activation supporting cognitive empathy, although this effect did not reach significance (B = 0.17, β = 0.20, t(86) = 1.89, p = 0.061) (Fig. 5A). We additionally tested for the difference in slopes for the regression, with an interaction term between age as a continuous variable and condition (affective and cognitive empathy). We found a significant interaction between age and condition on brain activation, both when considering the HbO_2_ (b = -0.34, t(173) = -2.74, p = 0.007) and the HHB signal (b = -0.27, t(173) = -2.85, p = 0.005) for the left STG, indicating that the slope for age differed between the two conditions. These results suggest that the effect of age on brain activation varies depending on the condition (Fig. 5B).

**Figure 5.**
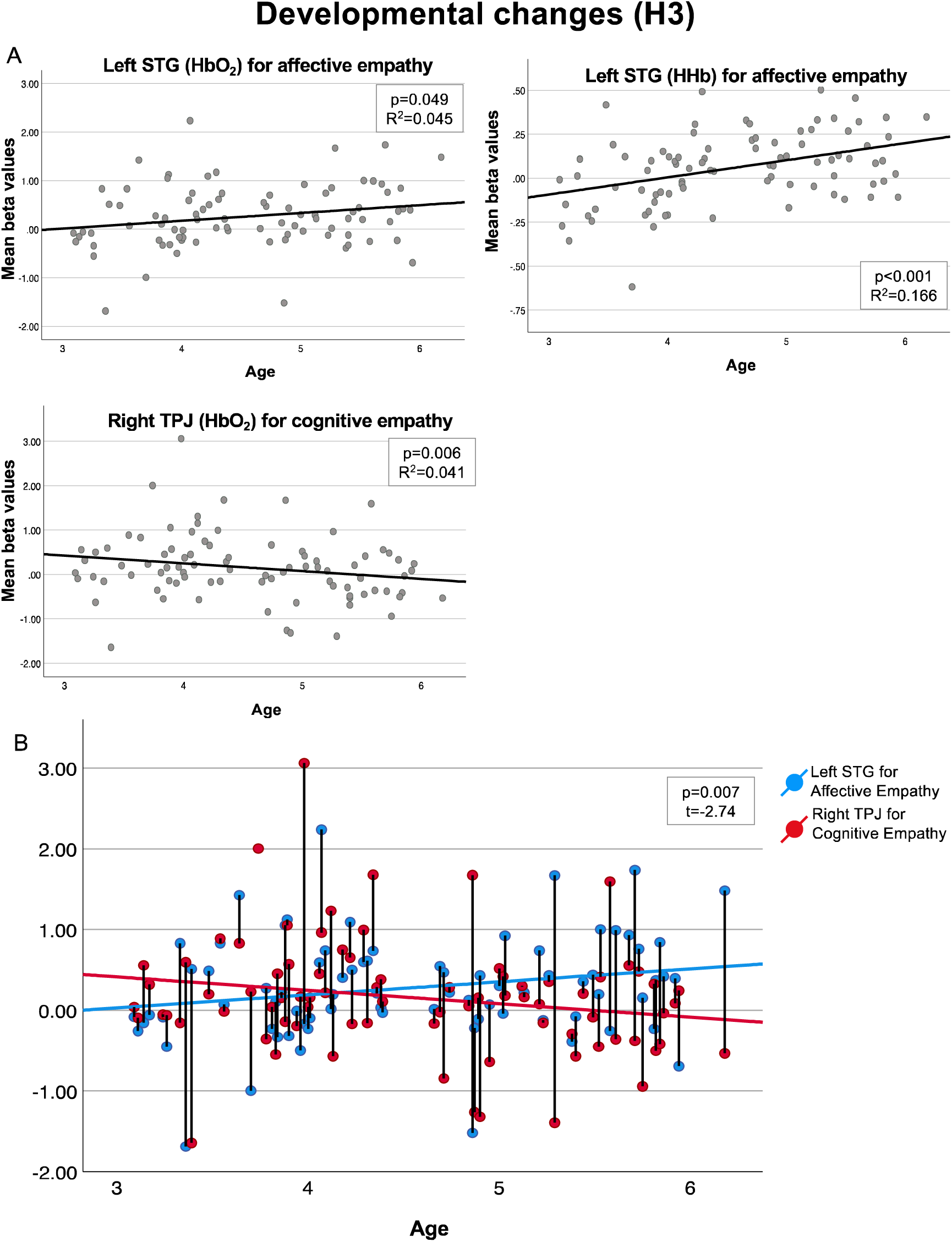
A) Scatterplots of the relationship between age as a continuous variable and mean beta values of left STG for affective empathy (both HbO_2_ and HHb) and right TPJ for cognitive empathy. The black line represents the regression line of best fit. B) Individual beta values of left STG (HbO_2_) for affective empathy and right TPJ (HbO_2_) for cognitive empathy across ages. The blue and red lines represent the line of best fit for affective and cognitive empathy. The vertical black lines join the two data points from the same participant.

To understand further the effect of age on the brain activations underlying affective and cognitive empathy, we performed one-way ANOVAs on the regions that showed significant different activations in the two empathy components (affective empathy, i.e. emotional events > neutral events, and cognitive empathy, i.e. questions on emotional state > questions on neutral facts) as dependent variables, and younger and older preschoolers (based on median split at age 4.35 years, N=45 per age group) as a between-factor. We found a significant effect of age group on the greater activation of the left STG (HHb) for emotional versus neutral events (*F*(1, 86) = 11.90, *p* < 0.001, η = 0.005), with older participants showing greater activation supporting affective empathy than younger participants (*t*(85) = 3.45, *p* < 0.001). Since we had hypotheses regarding the engagement of the MPFC in affective empathy, we additionally conducted a one-way ANOVA on MPFC activation for affective empathy. This was done to account for the possibility that if age plays a significant role in the activation of this region (given the later maturation of prefrontal cortex (43)), its activation may not be significant when collapsed across age groups. Indeed, we found a significant effect of age group on the greater activation of the right MPFC (HbO_2_) for emotional versus neutral events (*F*(1, 79) = 4.85, *p* < 0.031, η = 0.005), with older participants showing greater activation supporting affective empathy than younger participants (*t*(78) = 2.21, *p* = 0.015). We also found a significant effect of age group on the greater activation of the right TPJ (HbO_2_) for emotional vs neutral questions (*F*(1, 86) = 7.27, *p* = 0.008, η = 0.079), with younger participants showing greater activation supporting cognitive empathy than older participants (*t*(85) = 2.69*, p* = 0.004). Confirming that age-related change differed between cognitive and affective empathy, a repeated measure ANOVA on the preschoolers’ significant brain activations for the two empathy components (left STG for affective empathy vs right TPJ for cognitive empathy) as within-participant factors and age group as a between-factor (younger vs older preschoolers based on a median split) showed a significant interaction between condition and age group (HbO_2_: *F*(1, 84) = 4.94, *p* = 0.029, η^2^ = 0.056; HHb: *F*(1, 84) = 11.37, *p* = 0.001, η = 0.119).

Results from all these analyses converge on identifying an effect of age on the neural underpinnings of affective and cognitive empathy, with younger preschoolers showing greater activation in regions supporting cognitive rather than affective empathy, and older preschoolers showing greater activation in regions supporting affective rather than cognitive empathy.

## Discussion

Understanding whether affective or cognitive empathy emerge in early development is of paramount importance in developmental psychology, as this sheds light on the mechanisms underlying the emergence of empathy and, more importantly, informs how we can foster the development of this social ability. While the field so far has widely assumed that cognitive empathy arises from the roots of the affective component, we challenged this idea by suggesting that understanding others’ emotions might precede feeling them. Importantly, this work provides the first empirical evaluation for our proposed *foundational cognitive empathy model* (19), showing that the specialisation of brain networks supporting empathy components changes with age – i.e. with younger preschoolers’ brains exhibiting greater activation for cognitive empathy, while older preschoolers’ brains responding more to affective empathy. If replicated, these results will be crucial to suggest a new framework for understanding the development of empathy.

### Brain regions supporting empathy in young children resemble the adult ones (H1)

Examining neural and physiological markers of empathy provided a crucial window into the dynamics of empathy development, allowing us to not solely relying on its behavioural and verbal manifestations that might be limited in young children. Building upon the extensive neuroimaging literature in adults identifying the brain regions supporting empathy, we first aimed to explore whether the same brain regions supporting affective and cognitive empathy in preschoolers as well. Indeed, we found clear network specialisation for these two empathic components even at preschool age, with the left STG supporting affective empathy and the bilateral DLPFC, right MPFC and TPJ supporting cognitive empathy.

The superior part of the temporal lobe has been extensively documented as a crucial hub for social perception (44–46), particularly in processing information from faces, such as facial expressions (47, 48). Additionally, the role of the STG in sharing others’ emotions has been widely documented, particularly for its key role within the Mirror Neuron System (MNS) (49, 50). In this context, and for its role in imitation, this region is thought to support the simulation of sensory consequences of the observed action (51–53). Additionally, the STG has strong anatomical and functional connections with subcortical limbic areas, such as the insula and amygdala, which are well-known for their involvement in processing emotional stimuli and in sharing others’ emotions (14, 53, 54). Unfortunately, due to the intrinsic limitations of fNIRS, we were unable to record activity from these subcortical regions. Future studies, however, could extend this work by recording from other cortical regions of the MNS, such as the inferior frontal gyrus, which were not recorded here due to a limited number of optodes. This would help confirming the involvement of the MNS in supporting affective empathy in young children. Interestingly, we did not find MPFC activation for affective empathy at the group level. However, consistent with our hypotheses, we found an effect of age on the activation of the right MPFC for affective empathy, with older preschoolers showing significantly greater activation than younger ones. This suggests that the non-significant results at the group level may have been driven by the younger participants, and that the engagement of this region in affective empathy emerges only later in development, which is consistent with our hypothesis and theoretical framework.

Cognitive empathy supported by the DLPFC and TPJ largely overlap with findings from adult studies (8–10). Interestingly, we also found engagement of the medial portion of the PFC for cognitive empathy. This finding has also been documented in some studies on adults (55), and it is consistent with the functional overlap between the affective and cognitive sides of empathy(15, 16). This engagement might be interpreted in light of the key role of the MPFC in self-reflection and self-processing (which we recently confirmed in the developing brain too (56)), a fundamental process for reflecting on others’ mental state and linking them to one’s own (57).

Interestingly, we did not observe any greater brain activations for neutral events compared to emotional events, or for questions about neutral states compared to questions about emotional states. Together with the significant emotional > neutral activations reported above, this indicates that condition differences were specific to emotional processing rather than reflecting general differences in task difficulty or attentional demands. Moreover, this indicates that, even from the age of three, children’s brain regions are more engaged by emotional events and reasoning than by events and reasoning lacking emotional connotations.

### Greater physiological response for negative compared to positive emotional events, which are also supported by specific brain areas (H2)

Our second prediction was that we would find greater reactions to negative compared to positive emotional events, aligning with the ’negativity bias’ hypothesis (27). Our heart rate results strongly support this, showing a robust and statistically significant decrease in heart rate specifically for negative emotional events, compared to positive and neutral ones. Although we did not observe greater brain activation exclusively for negative compared to positive emotional events, we identified preschoolers’ brain regions that selectively respond to different emotional valences, with the temporal lobe showing greater engagement for negative emotional events, and the bilateral MPFC for positive scenarios. While comparisons of brain regions supporting events with different emotional valences are quite limited in adults, consistent with our findings, the engagement of the MPFC for positive compared to negative events has also been documented in one adult study (58). This may suggest that the ‘negativity bias’ may be valid particularly at the physiological level, but at the neural level, children as young as 3 years of age are able to discriminate and react differently for positive and negative emotional events. This would be consistent with evidence on adults showing that different brain networks are engaged for pleasant and unpleasant stimulation (59). Our results show that this cortical specialisation for stimuli of different emotional valence, well documented in infancy and early childhood (60) is also engaged in the context of an empathy task in preschoolers, to our knowledge the first demonstration of valence-specific cortical responses within an empathy paradigm at this age. We hypothesized that this ’negativity bias’ would be stronger in younger compared to older participants, but did not find statistical evidence in support of this.

We found positive associations between MPFC activation for positive compared to negative events and greater heart rate decrease for positive events only. Why these associations were found only for positive and not negative emotional events is unclear, but one hypothesis is that preschoolers who may have engaged in more self-reflections during positive rather than negative emotional scenarios are those who showed a greater decrease in heart rate during emotional events. Physiological reactions to negative emotional events are more pervasive and less variable, which may explain why they were not statistically associated with significant brain activations during emotional events.

### Stronger markers of cognitive compared to affective empathy in younger preschoolers (H3)

In this work, consistent with our hypotheses and theoretical framework (19), we presented evidence supporting our proposed *foundational cognitive empathy model*, with regions supporting affective empathy more engaged in older preschoolers and regions supporting cognitive empathy more engaged in younger preschoolers. We interpret this as challenging years of speculation about the pivotal role of affective empathy in the development of its cognitive dimension, which was nevertheless supported by limited empirical evidence and not robustly investigated. We note, however, that a longitudinal design would be required to build on the current cross-sectional study of 3-to-6-year-olds to precisely establish the order in which these components first emerge within a child. Importantly, the age-related differences we observed were specific to the empathy-relevant contrasts rather than to overall task engagement. Because our design compared emotional versus neutral events, and questions about emotional states compared to questions requiring cognitive reasoning, using the same stimuli within the same children, generic social, attentional, and linguistic demands were present across conditions and are therefore unlikely to account for the effects. While we cannot fully exclude a contribution from the maturation of more general processes recruited by the task, an account based on these specific empathy-relevant distinctions is more parsimonious than one invoking generic developmental change, since it was these contrasts, not overall activation, that varied with age.

One could argue that the fact that younger preschoolers showed greater activation in regions supporting cognitive rather than affective empathy does not necessarily mean that young children did not engage *any* brain regions for affective empathy, and that the greater engagement of cognitive empathy regions might reflect that the younger portion of our sample struggle more to answer questions about others’ emotional states compared to older preschoolers. This interpretation aligns with the fact that younger preschoolers answered less congruently with the character’s emotions at the ‘*how does s/he feel?*’ question compared to older ones. However, there were no significant age differences for the ‘how *does this make you feel?*’ question, which arguable taps the cognitive reflection on one’s own emotional state rather than affective sharing per se. While accuracy on questions about others’ emotions and about neutral facts improved with age, accuracy on this self-reflective question did not, suggesting that age-related gains were specific to reasoning about others and general performance rather than to self-directed emotional reflection. Moreover, the number of participants who did not make *any* mistake was remarkably high regardless of age. It is important to note that during the testing sessions, we noticed that some participants (∼16%) deliberately and consistently responded in the opposite way to what was expected to the questions on their own or others’ emotional state (a typical challenging preschoolers’ behaviour!), and even smiled after the wrong answer or commented on this after the task. This underlines the necessity to rely not only on verbal or behavioural manifestations of empathy, but also on other objective measures, when investigating this skill at such a young age. Nonetheless, even if one believes that evidence of cognitive empathy in young preschoolers does not necessarily exclude the presence of affective empathy, this still suggests that preschoolers begin developing cognitive empathy as early as three years old, at least alongside the affective dimension.

Importantly, considering the cognitive dimension of empathy as foundational for the development of empathic reactions per se may also inform the debate on whether empathy is learned or innate (61). In fact, if understanding others’ emotions is crucial for sharing them, then anything that supports *this* understanding becomes crucial for an empathic sharing to arise. Our data suggest that younger preschoolers engage more when *explicitly* prompted, while older preschoolers may have learned to think about what others are experiencing. This is consistent with findings from theory of mind/mind-reading development (62), with recent evidence showing that different brain networks support the implicit or explicit versions of this skill (63). Moreover, it highlights the important role of parental scaffolding and verbal engagement in helping children develop empathy. It is in fact plausible to think that the cognitive dimension of empathy may not arise automatically, but it is prompted and fostered through others’ scaffolding. In this context, the role of significant adults – i.e. carers, parents, teachers – becomes pivotal in supporting the understanding of what others are feeling in young children. A recent meta-analysis has shown that parents’ mental state talk is positively associated with their children’s performance in false belief and emotion understanding in participants older than seven years of age (64); therefore, it is highly plausible that parenting and modelling positive behaviours will also promote empathy development in younger children. This would also be in line with Bandura’s social learning theory (65), which has received substantial empirical support for the development of social skills (66). Interestingly, this pattern might also be compatible with an earlier development of cognitive empathy over affective empathy from a phylogenetic perspective, where the ability to model and understand others’ emotions may have provided evolutionary survival advantages even before emotional resonance developed. Therefore, from an evolutionary standpoint, navigating complex social situations may have been as important, if not more so, than fostering emotional bonding (67, 68).

This groundbreaking empirical evidence possibly supporting of our proposed *foundational cognitive empathy model* may have significant implications for educational practices in the early years, opening up new possibilities for behavioural interventions that are often dismissed or applied too late, as children with low-empathic traits or poor social skills are often perceived as ’natively bad’. While the Early Years Foundation Stage (EYFS) framework (adopted in UK classrooms for children aged five to seven (69)) recognises ‘Personal, Social, and Emotional Development’ as a key focus for early years (70), most of the suggested classroom activities are not research-based. Importantly, this work is one of the few studies to investigate markers of empathy in preschoolers, and the only to focus on specific dimensions of empathy at such young age, enabling the design of interventions to foster this skill in educational settings early in development, possibly before dysfunctional patterns of empathic responses are established. Results of this work could pave the way for a new line of research in educational neuroscience, which could inform EYFS practices, especially regarding the importance of modelling empathy during class activities. This could have a cascade effect into government policies, as children with poor social skills currently represent a substantial high cost for the society and for their families (71, 72).

### Strengths and limitations of this work, and future directions

The robustness of our results relies on a large sample size (exceeding the preregistered one), which is particularly noteworthy for an fNIRS study with developmental populations. Our statistical analyses were preregistered, strongly hypothesis-driven and motivated by a theoretical framework we previously shared with the scientific community (73), and all analyses examining brain region activations under different conditions were corrected for multiple comparisons.

The contribution of this work to advancing the field of developmental neuroscience in understanding empathy development would not have been possible without designing a novel task specifically suited to assess empathy components in preschoolers, following a thorough review of the limitations of previously used tasks in the literature. This task can be shared with the scientific community upon request to the corresponding author and potentially used with children of different ages and backgrounds. It could furthermore be implemented in functional MRI studies to assess the role of subcortical structures in older children. Our low attrition rate for both neural and physiological data highlights the strong feasibility of this task in capturing markers of empathy beyond behavioural and verbal manifestations. Furthermore, the remarkably high percentage of participants who answered all questions correctly or as expected confirms that the scenarios were well-suited for this age group. Building on this, future work should validate the scenarios in independent and more diverse populations, including children from different cultural and linguistic backgrounds and across a broader age range, to confirm that the stimuli are interpreted as intended and to establish the generalisability of the paradigm.

We acknowledge that the characters in the scenarios all displayed white Caucasian features, which matched the ethnicities of most of our participants. All preschoolers tested attended daycare in London, so also those participants who might have not identified as white were exposed daily to children from diverse cultural and ethnic backgrounds, including white Caucasian. However, this could be considered a limitation if the task is used with children from predominantly different ethnic backgrounds or those less exposed to racial diversity in the future. In this work, although we initially planned to design characters with different ethnic and racial features for the scenarios, we later realised that randomising all possible combinations of main character/s exposed to emotional or neutral events vs other characters in the scene either matching or not matching the participant’s ethnicity and race would have resulted in a high number of ingroup/outgroup combinations that would have not been feasible within just the 16 trials planned. We therefore chose to use characters from a single ethnic background and expose all participants to the same set of stimuli.

The results from this work open up new interesting avenues for research, as future studies should investigate which component of empathy is selectively impaired in children with developmental conditions, in order to inform targeted interventions. For instance, some have suggested that children with autistic traits are capable of sharing others’ feelings but struggle with the more cognitive aspects of empathy (74). On the other hand, it has been proposed that children with high callous-unemotional traits, characterised by reduced affective empathy alongside relatively intact cognitive empathy, may be able to understand others’ emotional states without sharing them (75, 76), which may enable them to manipulate others without remorse. In these children, whether the affective component of empathy develops first and then decline or whether just the cognitive one develops is still debated (77). However, more recent evidence suggests that both children with high callous-unemotional traits and autistic children may be deficient in both dimensions of empathy (78, 79). As much of the current knowledge on empathy in children with developmental conditions comes from studies using the limited tasks described in the introduction, and the field has yet to reach a clear consensus on whether both components of empathy or just one are impaired in children with social difficulties, employing our task to explore the neural and physiological underpinnings of empathy could provide a more robust understanding of the developmental trajectories of these empathic components in such children, so to design more tailored interventions.

## Conclusions

In this work, we showed that brain regions supporting affective and cognitive empathy in preschoolers resembles the adult ones, suggesting a network specialisation from very early on in life. More importantly, we challenged existing views providing the first evidence that cognitive empathy might arise earlier than the affective dimension. There results hold paramount importance on understanding social development, and have implications for targeting interventions, and for considering the pivotal role that adults might have in scaffolding the understanding of others’ emotions.

## Materials and Methods

This study was preregistered on the Open Science Framework website (https://osf.io/crvz2).

### Participants

Ninety-five 3-to-6-year-olds observed the stimuli (53 males, age mean ± SD = 4.52 ± 0.84 years) while we simultaneously measured their heart rate and their neural responses using fNIRS. Looking time patterns and pupil size was also recorded with the Eyelink Eyetracker but these data are not presented in this work. Seven additional participants were recruited but refused to wear the equipment or did not do the task. Out of the ninety-five participants assessed with this task, 3 were excluded because of: (i) technical issue (1 participant); (ii) experimental error (2 participants). For the fNIRS analyses, we additionally excluded 2 participants for poor data quality. As a result, ninety participants were included in the fNIRS analyses (50 males, age mean ± SD = 4.53 ± 0.84 years). For the heart rate analyses, we additionally excluded 2 participants because they did not want to wear the electrocardiogram (ECG) electrodes, and 2 participants for technical issues with the ECG system. As a result, ninety-one participants were included in the heart rate analyses, (50 males, age mean ± SD = 4.52 ± 0.85 years). A total of 87 participants were included in both the fNIRS and the heart rate analyses (our target sample was 75 participants, as preregistered here https://osf.io/crvz2/overview).

All included participants were born full-term, healthy and with normal birth weight. Participants were excluded from recruitment if they had a known significant neurodevelopmental condition or a medical condition that was likely to impact brain development or impede the child’s ability to participate in this study. Written informed consent was obtained from the toddler’s caregiver prior to the start of the experiment. Ethical approval for this study was given by the Ethics Committee of the Department of Psychological Sciences, Birkbeck, University of London (No. 2122056).

### Stimuli and procedure

Participants were tested in a sound attenuated room, sitting on a chair at approximately 60 cm from a plasma screen. Participants were presented with a standard block design task with 8 emotional and 8 neutral scenarios. Among the 8 emotional scenarios, 4 were negatively connoted and 4 were positively connoted. The order of the conditions was pseudorandomized to ensure that participants were not exposed to the same condition more than twice in a row. A new pseudorandomized order was generated for each participant. Scenarios were alternated with 9-12 seconds jittered baseline, showing swirly coloured images. The experiment ended if the participant saw all the 16 scenarios or if s/he showed any sign of fussiness or discomfort.

Each scenario was composed of 3 pictures drew by a professional designer, with the first 2 displayed for 6 seconds (set-up) and the last one showed for 8 seconds (affective empathy or neutral fact). The presentation of each picture was accompanied by a voice of a native English speaker describing the scene. The third picture was followed by 8 seconds of baseline showing swirly coloured images. After the baseline, in emotional scenarios, the narrating voice first asked the participant whether the character in the scenario was happy or sad (cognitive empathy). After 8 seconds, the narrating voice asked whether this made the participant feel happy or sad (reflecting on their own feelings). In neutral scenarios, the narrating voice asked if the character in the scenario acted for one reason or another (cognitive reasoning) (Fig 1A).

The scenarios depicted situations that young children are typically exposed to, primarily involving children of their age. Specifically, the scenarios were drawn from existing studies on empathy in toddlers and children or were explicitly designed based on real-world episodes observed during 21 hours of nursery classroom observation conducted by our team (73). For the neutral scenarios, we acknowledge that scenarios depicting everyday activities cannot be entirely free of incidental affective associations; what defined a scenario as neutral was the non-emotional nature of its central event and the factual focus of its question.

### fNIRS data acquisition and array configuration

fNIRS data were acquired using two mobile systems (dual Brite MKII, Artinis Medical Systems BV, Netherlands) with a 48-channel array configuration as used in a previous recent work from our team

(80). Here the channels were coregistered onto an age appropriate MRI template from the Neurodevelopmental MRI Database of the University of South Carolina (http://jerlab.psych.sc.edu/NeurodevelopmentalMRIDatabase/)(81) to create ten regions of interest (ROI, left and right medial prefrontal cortex, MPFC; left and right dorsolateral prefrontal cortex, DLPFC; left and right temporo-parietal junction, TPJ; left and right middle and superior temporal gyrus, M/STG; left and right inferior parietal lobule, IPL) (see Table 1 and Figure 4 in 79).

### fNIRS data processing

Data analyses were carried out using in-house codes developed in MATLAB (MathWorks, Natick, MA) based on functions from Homer2 toolbox (82) and from the Statistical Parametric Mapping (SPM)–NIRS toolbox (83). Raw intensity data were converted to optical density (*hmrIntensity2OD.m* function from Homer2). Hereafter motion artefacts were corrected using a wavelet filter (iqr = 0.8, *hmrMotionCorrectWavelet*.m function from Homer2). We did not correct for motion artifacts using also spline as originally planned in the pre-registration, as the fNIRS data quality was overall good, considering that preschoolers were sitting and watching a screen during the task and were not required to move. We therefore chose to alter the data as little as possible. Low-quality channels were pruned based on physiological indicators of quality using QT-NIRS (https://github.com/lpollonini/qt-nirs). The software takes advantage of the fact that the cardiac pulsation is recorded in addition to the haemodynamic activity (84), and quantifies its strength in the spectral and temporal domains with two measures: the Scalp Coupling Index (SCI) and the Peak Spectral Power (PSP) (85). For each participant, data quality was assessed channel-by-channel and SCI and PSP were calculated every 3-second in non-overlapping window (threshold SCI = 0.70, threshold PSP = 0.07, empirically defined). Channels that had both SCI and PSP below threshold for more than 60% of the windows, were excluded from further analyses. The surviving channels underwent visual inspection, and channels with clear signs of noise or saturation were additionally removed. Optical density data were then bandpass filtered (0.01–0.5Hz) (*hmrBandpassFilt.m* function from Homer2) and converted into relative concentrations of haemoglobin using the modified Beer– Lambert law (DPF = 5.4, 4.6) (86) (*hmrOD2Conc.m* function from Homer2).

For each participant, we then constructed a design matrix with twelve regressors: i) baseline trials; ii) event for the emotional negative scenarios; iii) events for the emotional positive scenarios; iv) events for the neutral scenarios (8 second picture); v) first questions for the emotional negative scenarios; vi) first questions for the emotional positive scenarios; vii) questions for the neutral scenarios; viii) second questions for the emotional negative scenarios; ix) second questions for the emotional positive scenarios; x) baseline trials in between the events and the questions; xi) set-up scenes (12 seconds before the event, i.e. the first two 6 seconds pictures); xii) averaged signal of the two short-separation channels (one over the frontal and one over the temporal cortex) to account for systemic changes (87, 88). The regressors were convolved with the standard hemodynamic response function (HRF) t to generate the design matrix (89). This design matrix was then fit to the data using the general linear model (GLM) as implemented in the SPM–NIRS toolbox. Beta estimates were obtained for each participant for each of the regressors. The betas were then used to compute the following contrasts and test our hypotheses through one sample t-tests:

- affective empathy: (events for the emotional negative scenarios + events for the emotional positive scenarios) > (events for the neutral scenarios), computed over the 8-second event phase;

- cognitive empathy: (first questions for the emotional negative scenarios + first questions for the emotional positive scenarios) > (questions for the neutral scenarios), computed over the 8-second event phase;

- emotional valence: (events for the emotional negative scenarios) > (events for the emotional positive scenarios), computed over the 8-second event phase. Fig 1A shows a graphical representation of the contrasts in the block design.

Note that the HbO_2_ signal was fit with the positive canonical HRF and the HHb signal with the negative canonical HRF, hence the direction of the t-values is the same for the two chromophores (i.e. a positive t-value/beta value represents an increase in HbO_2_ and a decrease in HHb). To ensure statistical reliability, we applied the false discovery rate (FDR) correction for multiple comparisons to the brain activations results (90). When an ROI had no channels surviving our quality criteria for a given participant, that participant was excluded from analyses involving that ROI. Therefore, degrees of freedom may vary across regions.

### Heart-rate data acquisition and data processing

Interbeat interval (IBI), the time between heart beats and related to a faster heart rate during the task was used as a physiological measure of arousal, likely linked to the experience of affective empathy (42). Heart rate data was collected using the RSPEC BioNomadix (BIOPAC Systems, Inc, Goleta, CA, USA), attached to a MP160 amplifier, sampled at 2000 Hz. Three ECG electrodes were placed in a lead-II position on the child’s chest. AcqKnowledge (version 5, BIOPAC Systems, Inc., Goleta, CA, USA) was used for data acquisition. QRS peaks were automatically identified in AcqKnowledge, followed by a visual inspection to detect any not identified or incorrectly identified peaks. After exporting the peaks in Matlab, data were segmented into two parts for each scenario: the setup phase (first two pictures of each scenario) and the event phase (the third picture of each scenario, corresponding to either the affective empathy event or a neutral fact). As previously done(91), missing peaks were corrected for up to three consecutively missing beats by dividing the interbeat interval. If more than three consecutive beats were missing, the segment was excluded from further analysis. For each participant, we calculated the average IBI for each part of the scenario (set-up or event) and then averaged these values for the three conditions (emotional negative, emotional positive, and neutral). To investigate whether there was a change in participants’ heart rate when attending the scenarios events, a repeated-measures ANOVA on the preschoolers’ average IBI with part of the scenario (set-up, event) and condition (emotional negative, emotional positive and neutral) as within-participant factors was performed.

## Acknowledgments

We are very grateful to all the preschoolers and parents who participated in this study. We thank Vanessa Madeira, Gila Ehrenstein and the placement and master students for research assistance during the testing sessions. We thank Karen Angelucci for drawing the pictures of the stimuli.

This work was supported by the Leverhulme Trust and undertaken at the Centre for Brain and Cognitive Development (CBCD), Birkbeck College, London. C.B. acknowledges support from the Leverhulme Trust Early Career Research Fellowship (ECF-2021-174). P.P. acknowledges support from the Wellcome Trust (212979/Z/18/Z). E.J.H.J. acknowledges support from the Economic and Social Research Council (ES/R009368/1) and from the Medical Research Council (MR/K021389/1; MR/T003057/1). T.B. acknowledges support from grant from the UK Medical Research Council (MR/X010716/1).

## Code and Data Availability

The data supporting the results of this paper and the task can be made available upon reasonable request to the corresponding author through a formal data sharing and project affiliation agreement.

